# Quantitative relationships for radiation induced chromosome instability: data analysis

**DOI:** 10.1101/224006

**Authors:** Y.A. Eidelman, S.V. Slanina, V.S. Pyatenko, I.K. Khvostunov, S.G. Andreev

## Abstract

The experimental observations demonstrate that different cell lines reveal various shape of dynamic curves for radiation-induced chromosomal instability (RICI). We analyzed our own and published data on RICI for three cell lines, CHO-K1, V79 and TK6, on the basis of the mechanistic RICI model. We demonstrate that all three dynamic curves can be successfully described by the proposed model with partially cell line specific parameters.

## INTRODUCTION

We have developed a computer model of radiation-induced chromosomal instability (RICI) [1]. This paper is devoted the analysis of the results of own experiments together with the published data [2, 3]. The findings and experimental procedure are presented in the following sections.

## METHODS

### Experimental design

The experimental study was performed with a continuous culture of CHO-K1 Chinese hamster cells (PanEco). Cells were seeded in 25 cm^2^ flasks with 10 ml of medium (density, 10^6^ cells per flask) and cultured for 1 day to obtain an exponentially growing culture. Cells were cultured at 37°C in a 5% CO_2_ atmosphere in DMEM (PanEco) supplemented with 10% cattle serum and antibiotics (penicillin 50 U/ml and streptomycin 50 mg/ml). Cells were γ-irradiated (^60^Co) at a dose rate of 0.25 Gy/min. After irradiation they continuously proliferated (subculturing with a density of 10^6^ cells per flask 3 days after the exposure (t_1_) and then every 2 days (t_2_, t_3_… in Fig. 1). Cytogenetic analysis was performed 1 and 2 days after irradiation and 1 day after each passage. The chromosome spreads were stained according to Giemsa. The chromosomal aberrations (dicentrics) in diploid cells were analyzed.

**Fig. 1.**
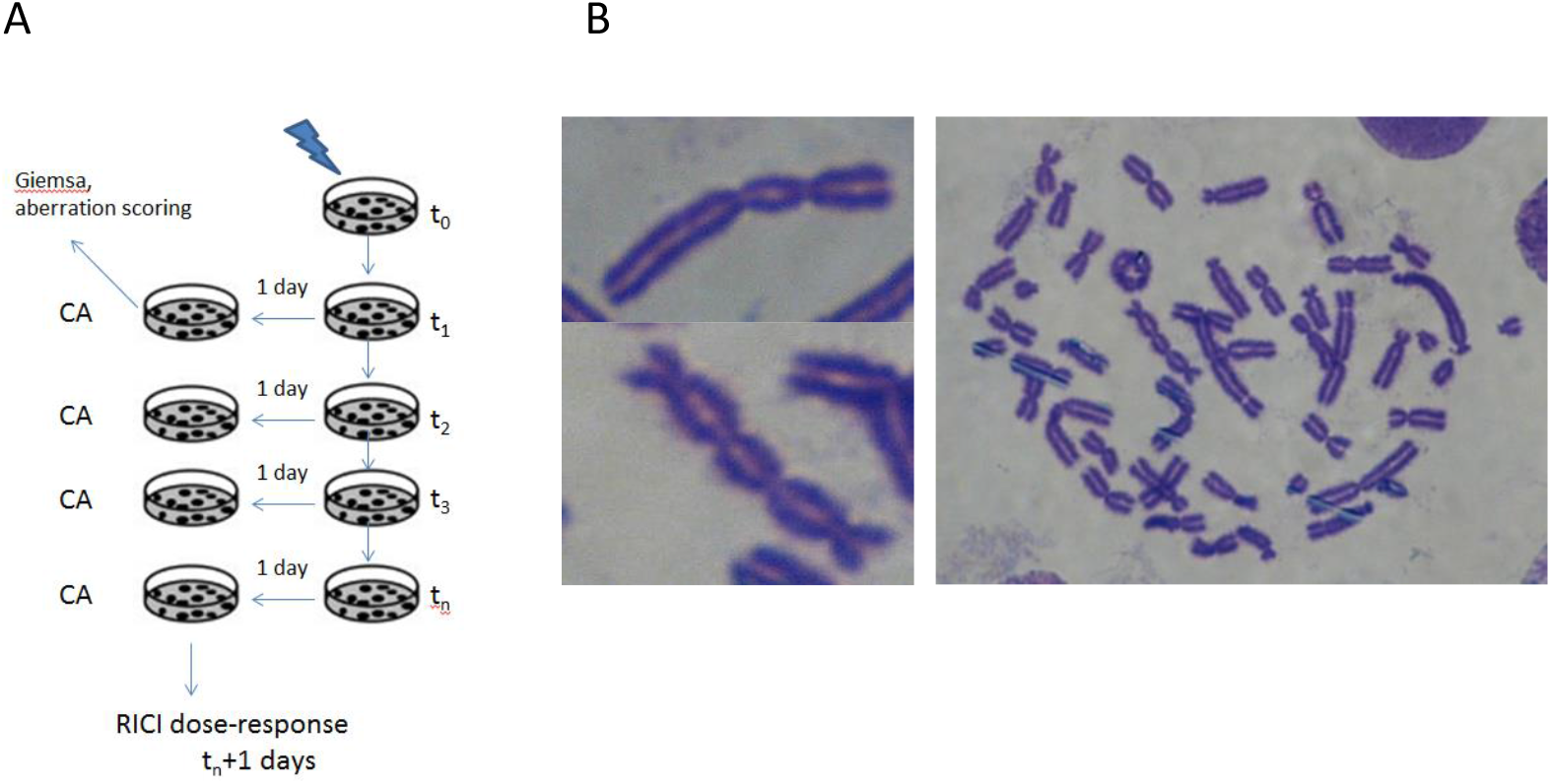
The experimental scheme used for RICI detection in CHO-K1 cells. A: the scheme of cell culturing, see Methods for explanation. B: photomicrography of chromosome aberrations (dicentrics), including complex aberrations, in CHO-K1 cells detected 6 days after exposure. The observed dicentrics and complex aberrations were classified routinely as follows: a quadricentric was counted as three simple dicentrics and a tricentric was counted as two simple dicentrics.

### Model

RICI modeling was carried out according to the scheme developed previously [1]. In brief, the model is implemented in five phases: induction of DNA DSBs in an asynchronous cell population by ionizing radiation; their misrepair and conversion to the aberrations of the chromosomal type in the first postirradiation cell cycle; formation of DSBs and their correct repair and misrepair, leading to aberrations in the progeny of irradiated cells; the passage of aberrant cells through mitosis; and the proliferation of the cell population. All of these processes, being stochastic, were modeled by Monte Carlo technique. The detailed model description can be found elsewhere [1].

## RESULTS

Fig. 2A shows the experimental data on the dynamics of dicentrics in CHO-K1 cells. The slow decline in the dynamic curve at a dose of 3 Gy was found. Notably, the experimental frequency does not reach the plateau at the maximum study time, 17 days post-irradiation. The experimental data and theoretical predictions were compared as shown in Fig 2B. The parameters of the model are presented in Table 1. The observed slow decline can be explained only if dicentrics are formed *de novo*. The reason is as follows. Supposing all the dicentrics observed in the progeny are radiation-induced dicentrics randomly transmitted to daughter cells in the first, second, and so on post-radiation mitoses, the decline of the slope of the frequency curve should be substantially sharper. Accordingly, dicentrics formed *de novo* are responsible for the difference between the red and the blue curves in Fig. 2B.

**Table 1.** The main parameters of RICI model for the presented theoretical curves.

| Cell line | CHO-K1 | V79 | TK6 |
| --- | --- | --- | --- |
| $P_s$ | 0.6 | 0.6 | 0.5 |
| $P_{bd}$ | 0.4 (1<sup>st</sup> 2 cycles), 0.25 (subsequent cycles) | 0.24 (1<sup>st</sup> cycle), 0.15 (subsequent cycles) | 0.2 (1<sup>st</sup> 2 cycles), 0.5 (subsequent cycles) |
| $P_{bi}$ | 0.5 | 0.6 | 1 |
| $P_{dbi}$ | 0.8 | 1 | 0.5 |
| $V$ | 0.0035 | 0.012 | 0.1 |
| $N_{dsb}$ | 0.2 | 2.1 | 2.6 |

$P_s$ – the probability of successful dicentric segregation;
$P_{bd}$ – the probability of death for a cell with an anaphase bridge ( $1 - P_{bd}$ – the cell survives, the bridge breaks);
$P_{bi}$ – the probability of broken bridge interaction with DSB;
$P_{dbi}$ – the probability of broken double bridge ends interaction with each other;
$V$ – chromatid dicentric formation rate, per DSB per hour;
$N_{dsb}$ – mean DSB generated per S phase, or converted from SSB during DNA replication.

**Fig. 2.**
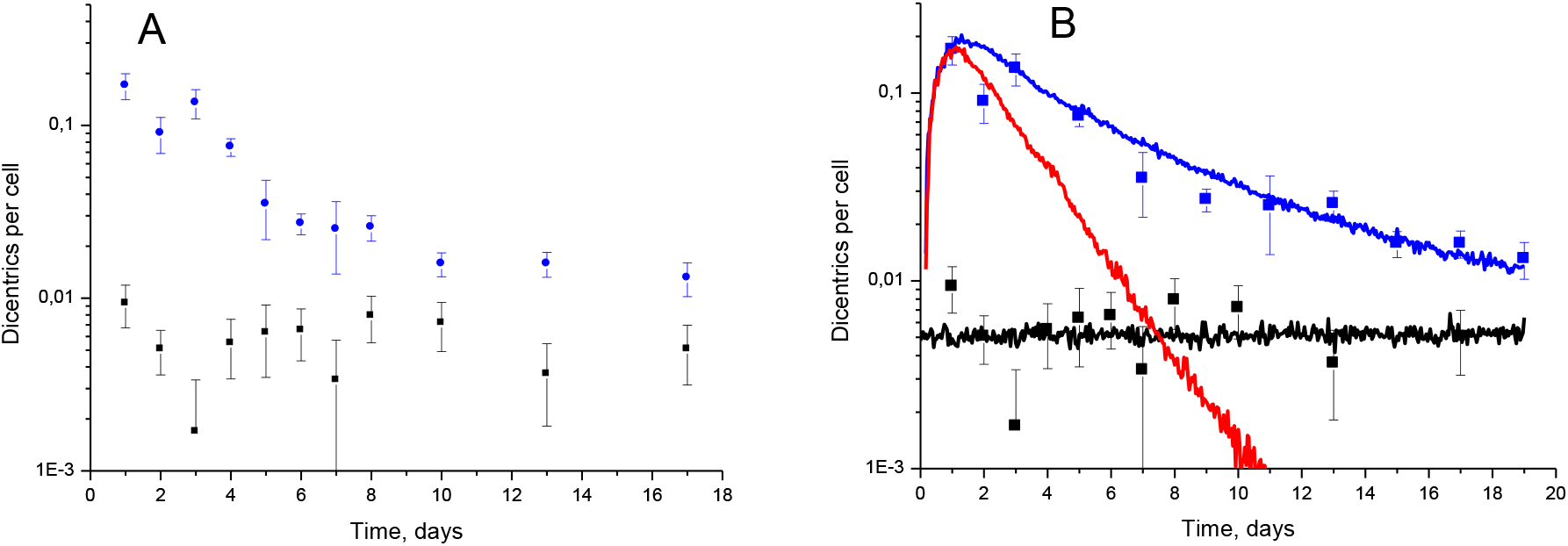
Dynamics of dicentrics after exposure, i.e. frequency of dicentrics as a function of time, measured in CHO-K1 cells. A – experimental data: control level of aberrations (black squares), 3 Gy (blue squares). B – data fitting using the theoretical model of RICI [1]. Black and blue curves correspond to control and 3 Gy exposure, respectively. The red curve is the calculated dynamics of dicentrics, provided that they appear in the progeny of irradiated cells only if cells have divided in mitosis correctly.

Fig. 3 shows the experimental data on the dynamics of RICI for the other two cell lines, V79 [2] and TK6 [3], together with their quantitative description by the theoretical model. It should be mentioned that the experimental data on TK6 cells [3] have been published dimensionless using FISH painting of four chromosomes. For comparability we merged and normalized the data so that the first point (t = 1 day) was close to the point corresponding to a dose of 3 Gy and a time of 30 hours in [4]. Two cell lines are characterized by different behavior of the dynamic curve: the V79 curve declines steadily and does not reach a plateau at the maximum examination time, 14 days, similar to CHO-K1 cells (Fig. 2). In the case of TK6 cells, the dynamic curve reaches a plateau at 10-15 days after exposure, and the height of the plateau is about ten times the control value [5]. Both types of dynamic curves, as well as the curve for CHO-K1, can be simulated in the framework of the same model using cell type-specific parameters. The main parameters for all types of cells are shown in Table 1.

**Fig. 3.**
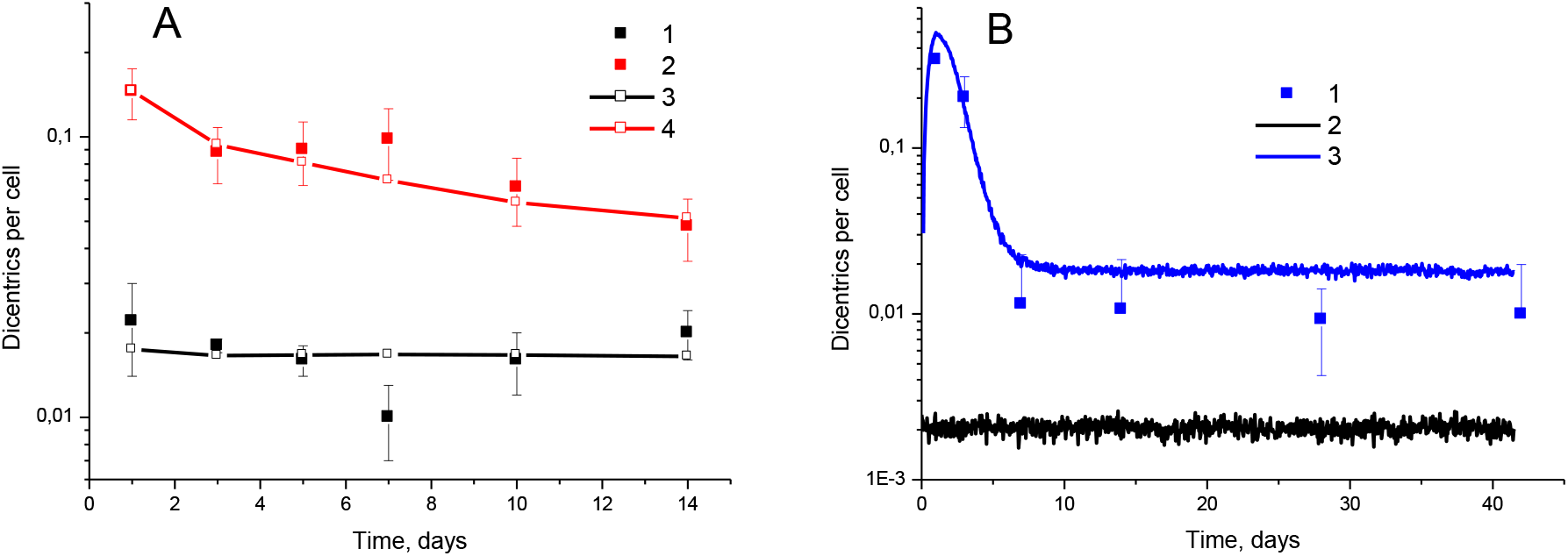
Dynamics of dicentrics in V79 and TK6 cells at different harvest times after exposure. A: V79 cells. 1, 2 – experiment [2]. 3, 4 – simulation. 1, 3 – control, 2, 4 – exposure with dose 3 Gy. B: TK6 cells. 1 – experiment [3], 3 Gy. 2, 3 – simulation. 2 – control, 3 – exposure with dose 3 Gy.

The comparison of the model curves calculated for different cell lines is shown in Fig. 4A. Fig. 4B shows the DSB generation rate for three cell lines obtained from the fitting of the analyzed data.

**Fig. 4.**
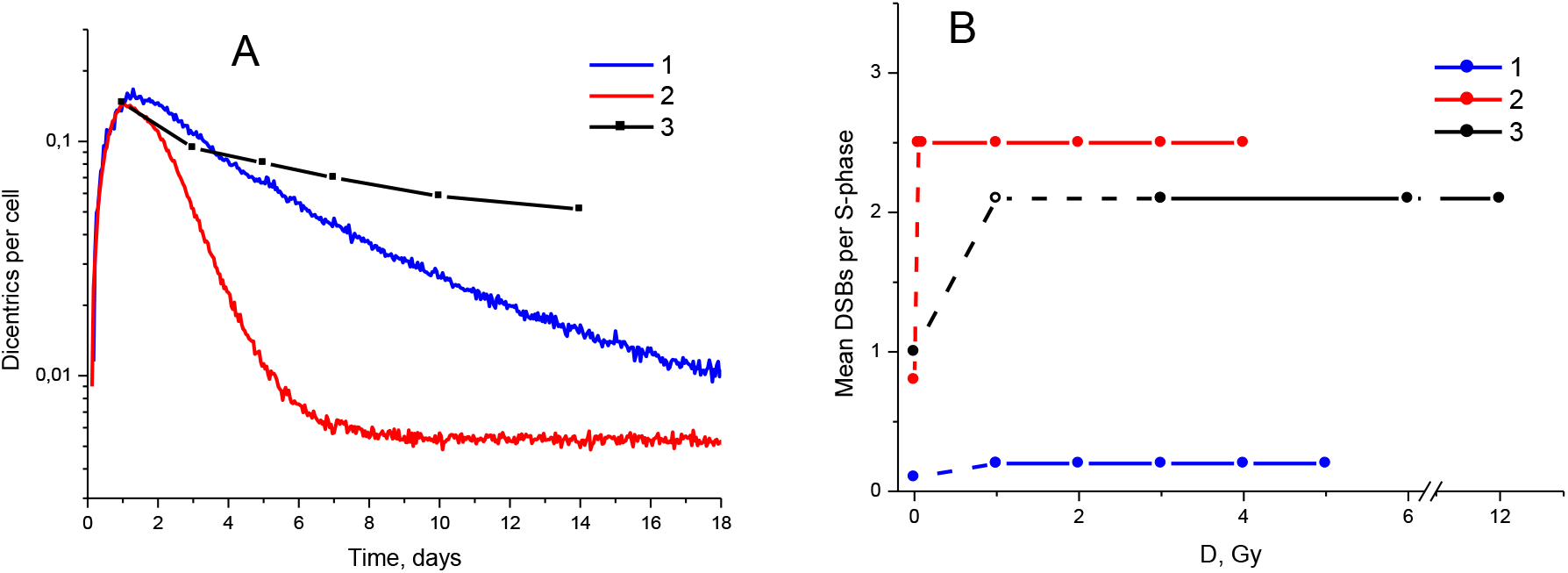
Comparison between theoretical curves of RICI for three cell lines: 1 – CHO-K1, 2 – TK6, 3 – V79. A: dynamics of dicentrics following 3 Gy irradiation. Theoretical curves are in accord with the experimental data shown in Fig. 2 and Fig. 3. All the curves are normalized to the point t = 1 day. B: theoretically reconstructed DSB generation rate in the progeny for different cell lines as a function of dose. DSB generation rate in the progeny of irradiated cells is constant, with the exception of the low-dose region, where it declines to the control level (unirradiated cells). The dashed lines and the hollow point are drawn in areas where no experimental data are available.

## DISCUSSION

The analysis of gamma-ray induced CI data (Fig 1–3, Table 1) revealed that the part of RICI mechanistic parameters are strongly cell type dependent and the other part is weakly dependent or independent. The agreement between theory and experiment in all cases was achieved on the basis of the general assumption that DSB generation rate in the S phase does not depend on a dose (in intermediate and high dose range) and time.

The assumption that DSBs can be spontaneously generated also in the G_1_ phase (in the progeny of irradiated cells) has to lead in appearance of dicentrics with accompanied paired chromosomal fragments. However this kind of dicentrics is not detected experimentally. Therefore this assumption was rejected, at least for the considered scheme of cell cultivation.

The developed RICI model was successfully applied to the interpretation of a set of experimental data on RICI dynamics. It should be noted that there are many unconfident data that were not included in analysis here. The probability of successful segregation of dicentrics (Fig. 1) was introduced as a new criterion to distinguish “novel” aberrations, appeared *de novo*, from “previous”, i.e. radiation-induced and just passively transferred from irradiated cells to their progeny.

The following two distinct types of dynamic curve shape were discovered in the course of RICI data analysis: one with a plateau and another one without a plateau, Fig. 4A. For the first time the set of RICI mechanism parameters which are inaccessible through the experimental evaluation were estimated by means of the theoretical RICI model, Table 1.

In conclusion, the developed RICI model successfully describes the *in vitro* data on the RICI dynamics (dicentrics) for different cell lines. The question of predicting RICI dose-dependencies on the basis of this model will be covered in the following papers.

## ACKNOWLEDGEMENTS

The present work was supported by Russian Foundation for Basic Research grant 14-01-00825 to S.A. S.A. acknowledges support from the MEPhI Academic Excellence Project (Contract No. 02.a03.21.0005).

## REFERENCES

1 Andreev S.G., Eidelman Y.A., Salnikov I.V., Slanina S.V. Modeling Study of Dose-Response Relationships for Radiation-Induced Chromosomal Instability. Dokl. Biochem. Biophys. 2013, 451, 171–175.

2 Jamali M., Trott K.R. Persistent increase in the rates of apoptosis and dicentric chromosomes in surviving V79 cells after X-irradiation. Int. J. Radiat. Biol. 1996, 70, 705–709.

3 Puerto S., Suralles J., Ramirez M.J., Creus A., Marcos R. Equal induction and persistence of chromosome aberrations involving chromosomes with heterogeneous lengths and gene densities. Cytogenet. Cell Genet. 1999, 87, 62–68.

4 Schwartz J.L., Jordan R. Selective elimination of human lymphoid cells with unstable chromosome aberrationsby p53-dependent apoptosis. Carcinogenesis 1997, 18, 201–205.

5 Schwartz J.L., Jordan R., Evans H.H., Lenarczyk M., Liber H. The TP53 dependence of radiation-induced chromosome instability in human lymphoblastoid cells. Radiat. Res. 2003, 159, 730–736.

6 Andreev S.G., Eidelman Y.A. Dose-response prediction for radiation-induced chromosomal instability. Radiat. Prot. Dosim. 2011, 143, 270–273.

